## Supplementary Materials for "Pericyte remodeling is deficient in the aged brain and contributes to impaired capillary flow and structure"

### Inventory of supporting information

**Supplementary Figure 1.** Comparison of basal vascular features in superficial cortex of adult and aged mice.

**Supplementary Figure 2.** Additional examples of triple pericyte ablations in adult and aged mice.

**Supplementary Figure 3.** Contributors to pericyte process growth potential.

**Supplementary Figure 4.** Pericyte processes distort capillary shape to gain endothelial coverage.

**Supplementary Figure 5.** Pericyte growth across microvascular zones in the adult and aged brain.

**Supplementary Figure 6.** Blood-brain barrier leakage is very rare with focal loss of pericyte coverage.

**Supplementary Figure 7.** No significant change in tight junction structure following the loss of individual pericytes.

**Supplementary Figure 8.** Apposition of astrocyte endfeet to the capillary wall is not overtly disrupted by focal pericyte loss.

**Supplementary Figure 9.** Dilations are localized to portions of the capillary segment lacking pericyte coverage.

**Supplementary Figure 10.** No dilations detected 3 days following off-target sham irradiation.

**Supplementary Figure 11.** Comparison of capillary diameters in awake versus isoflurane anesthesia conditions in aged mice.

**Supplementary Figure 12.** Capillary flow changes after focal capillary dilation in silico.

**Supplementary Figure 13.** Additional examples of capillary flow stalls and regressions.

**Supplementary Table 1.** Overview of selection criteria to choose potential base capillaries around which pericyte ablation will be mimicked.

**Supplementary Table 2.** Baseline characteristics for the set of affected capillaries around different base capillaries.

**Source data for all Main and Supplementary Figures.**

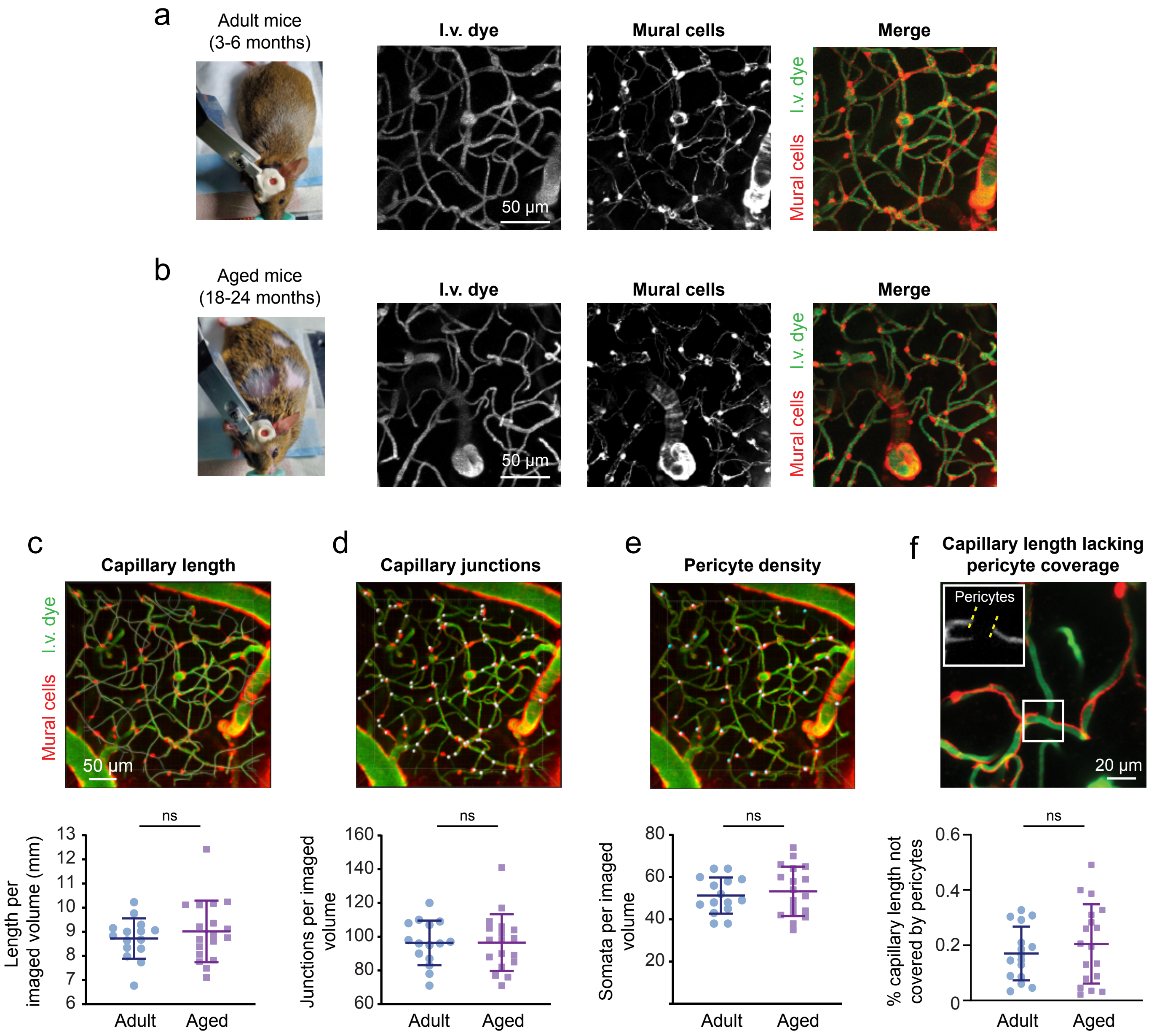

**Supplementary Fig. 1. Comparison of basal vascular features in superficial cortex of adult and aged mice. (a)** Representative image of a superficial capillary bed volume (10-100 µm from pial surface) in adult mice (3-6 months) and **(b)** in aged mice (18-24 months). I.v. dye = intravenous dye. **(c)** (Top) Representation of 3D reconstruction of the capillary vascular network. (Bottom) No difference in capillary length per imaging volume (350x350x90 µm regions) in adult versus aged animals, $t(31)=0.7711$, $p=0.4465$ by unpaired t test (two-sided). N= 15 regions from 9 adult mice, n=18 regions from 8 aged mice. **(d)** Example image of capillary junctions (white circles), and quantification showing no difference in capillary junctions per imaging volume (350x350x90 µm) in adult and aged mice, $t(31)=0.03123$, $p=0.9753$ by unpaired t test (two-sided) for n=15 volumes from 9 adult mice, n=18 volumes from 8 aged mice. **(e)** Example image of pericyte somata (cyan circles), and plot showing no difference in abundance per imaging volume (350x350x90 µm) in adult versus aged mice, $t(31)=0.5517$, $p=0.5851$ by unpaired t-test (two-sided) for n=15 volumes from 9 adult mice, n=18 volumes from 8 aged mice. Panels c,d,e are the same image. Panels c,d,e are the same image. **(f)** Exent of the capillary network not covered by pericytes, $t(32) = 0.8138$, $p=0.4216$ by unpaired t-test (two-sided) for n=15 volumes from 9 adult mice, n=18 volumes from 8 aged mice. Left image from panel (f) adapted from Berthiame et al. Frontiers in Aging Neuroscience, 2018 (PMID: 30065645). All data are shown as mean ± SD. All images representative of n=15 and n=18 volumes from adult and aged mice, respectively.

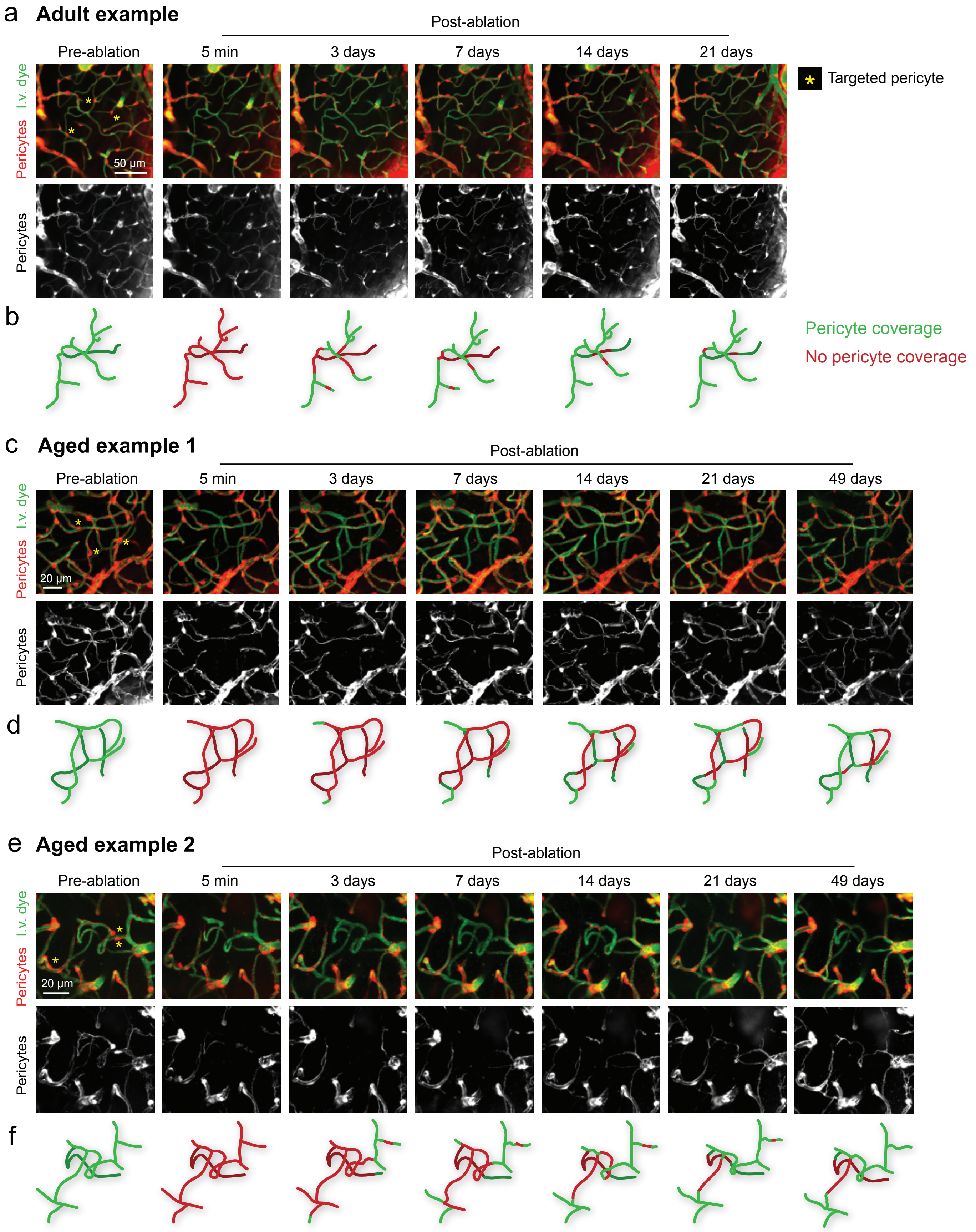

**Supplementary Fig. 2. Additional examples of triple pericyte ablations in adult and aged mice. (a)** Full time course of 21 day triple pericyte ablation in an adult mouse. Targeted pericytes marked with yellow asterisks. Bottom rows show pericytes only. Representative from 7 experiments in adult mice. I.v. dye = intravenous dye. **(b)** Schematic of capillary coverage over time. **(c)** Example of triple pericyte ablation experiment in an aged mouse, with additional chronic timepoint at 49 days post-ablation and **(d)** corresponding capillary coverage over time. **(e)** Second example of ablation from an aged mouse including 49-day timepoint. and **(f)** pericyte coverage of this capillary network over time. Panels c and e representative of 7 experiments in aged mice.

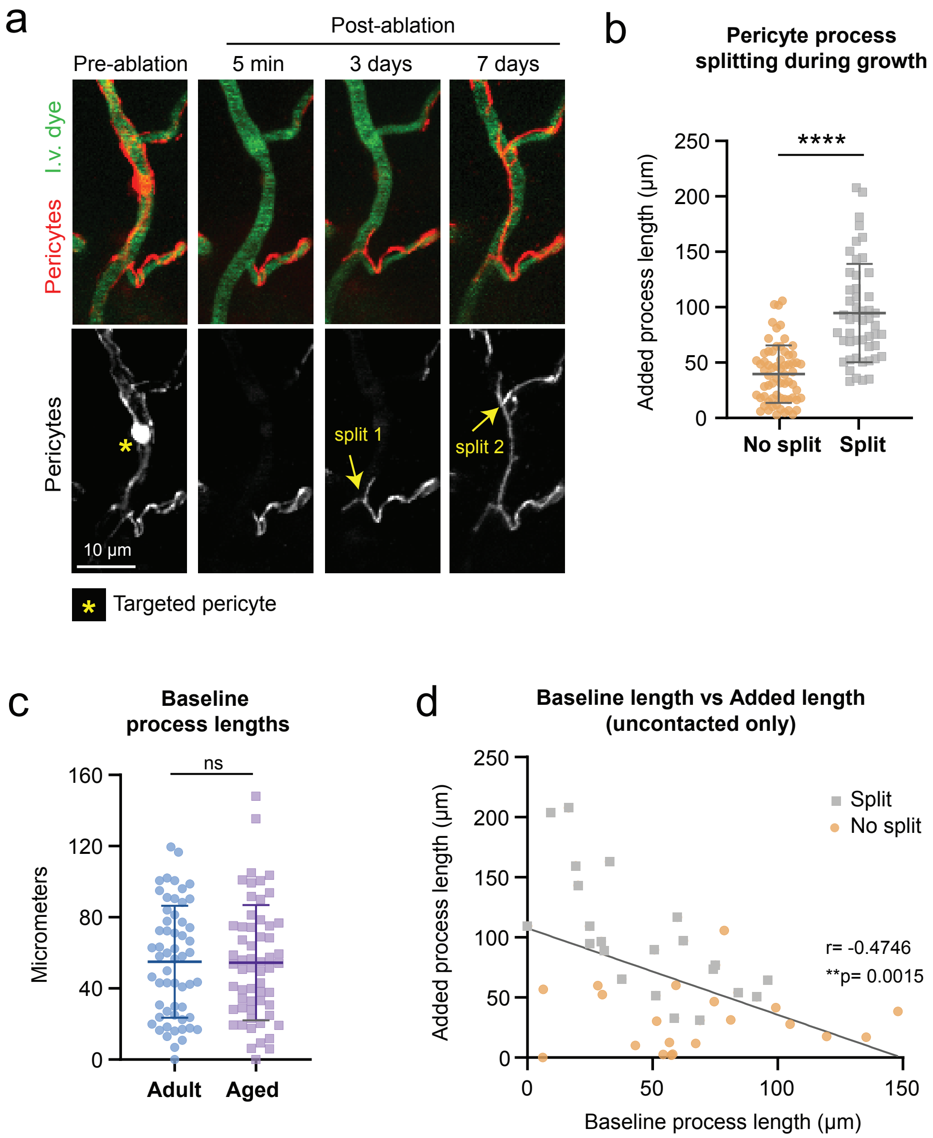

**Supplementary Fig. 3. Contributors to pericyte process growth potential. (a)** Pericyte process growing onto a capillary over time, with two occurrences of process splitting during extension. I.v. dye = intravenous dye. **(b)** Amount of pericyte process growth for processes that split during extension versus those that did not. T(73.79)=7.824, ****p<0.0001 by unpaired t test with Welch's correction (two-sided) for n=50 processes (no split), and n=66 processes (split) from 12 mice (6 adult, 6 aged). Presented as mean ± SD. **(c)** Baseline process lengths are no different across age groups. T(113)=0.0847, p=0.9327 by unpaired t test (two-sided). N=57 processes from 6 adult mice, n=58 processes from 6 aged mice. Data shown as mean ± SD. **(d)** Correlating baseline process length to length added during remodeling reveals a strong negative correlation. Processes that split versus those that did not are also noted. Pearson correlation (two-sided), r= -0.4746, F(1, 40)=11.63, **p=0.0015, for n=42 processes from 11 mice pooled across the two age groups (5 adult, 6 aged).

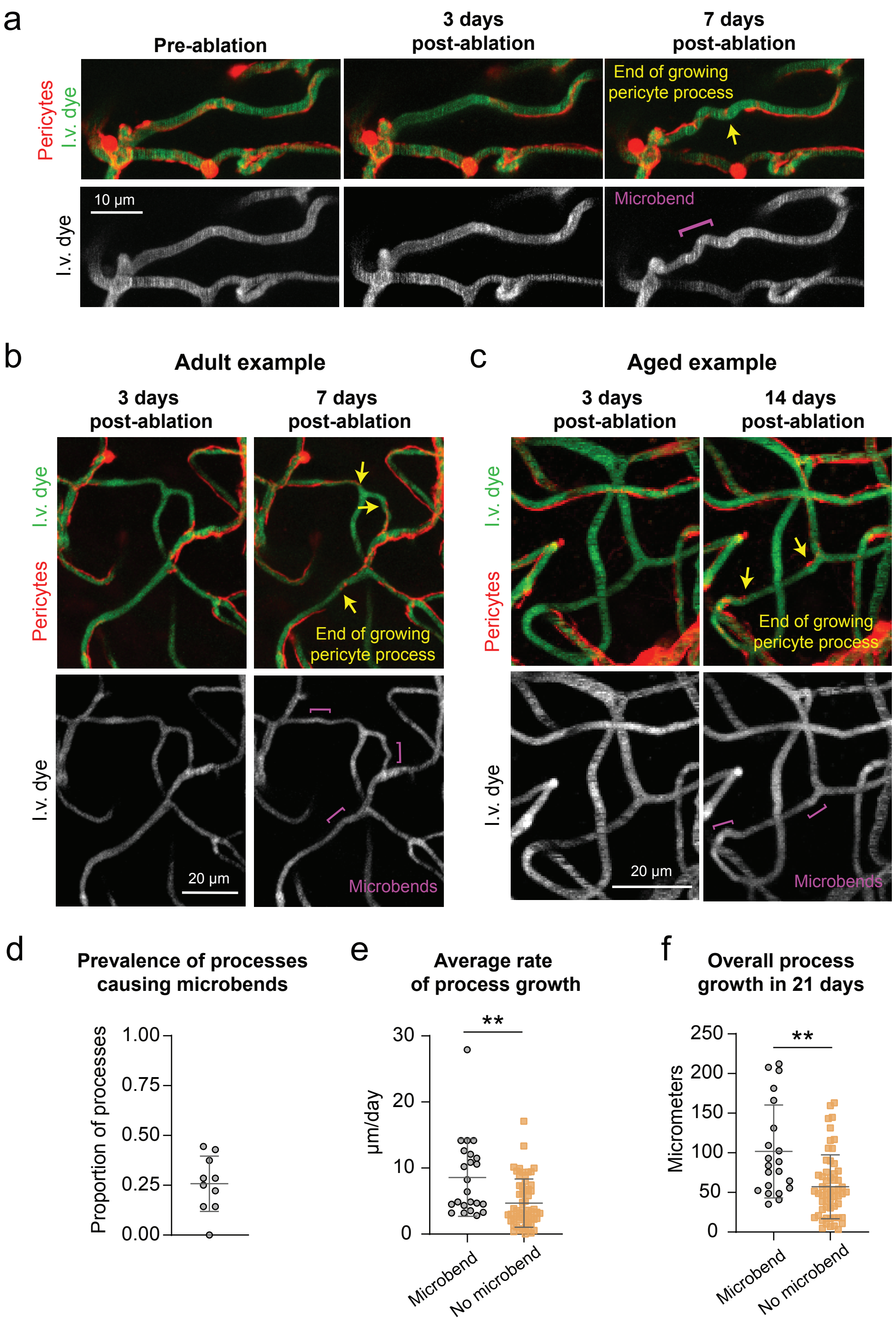

**Supplementary Fig. 4. Pericyte processes distort capillary shape to gain endothelial coverage. (a)** Tracking a remodeling pericyte growing over a capillary segment reveals the formation of a "microbend" near the growing process tip. I.v. dye = intravenous dye. **(b)** Examples of multiple microbends forming post-ablation in a triple ablation area in an adult mouse. **(c)** Multiple microbends created by remodeling processes in a triple ablation area in an aged mouse. **(d)** Plot of the proportion of processes which form a microbend in the underlying capillary during their extension. n=10 triple pericyte ablation experiments from 8 mice (4 adult, 4 aged). **(e)** Greater average rate of growth of processes that form microbends compared to those that do not. T(24.21)=2.922, **p=0.0074, by unpaired t test with Welch's correction (two-sided). **(f)** Greater overall final growth at Day 21 for processes that formed microbends compared to those that did not. T(25.54)=3.325, **p=0.0027 by unpaired t test with Welch's correction (two-sided). For both (e) and (f), n=20 pericyte processes creating microbends, 57 pericyte processes that don't create microbends, across 8 mice (4 adult, 4 aged). All data shown as mean ± SD.

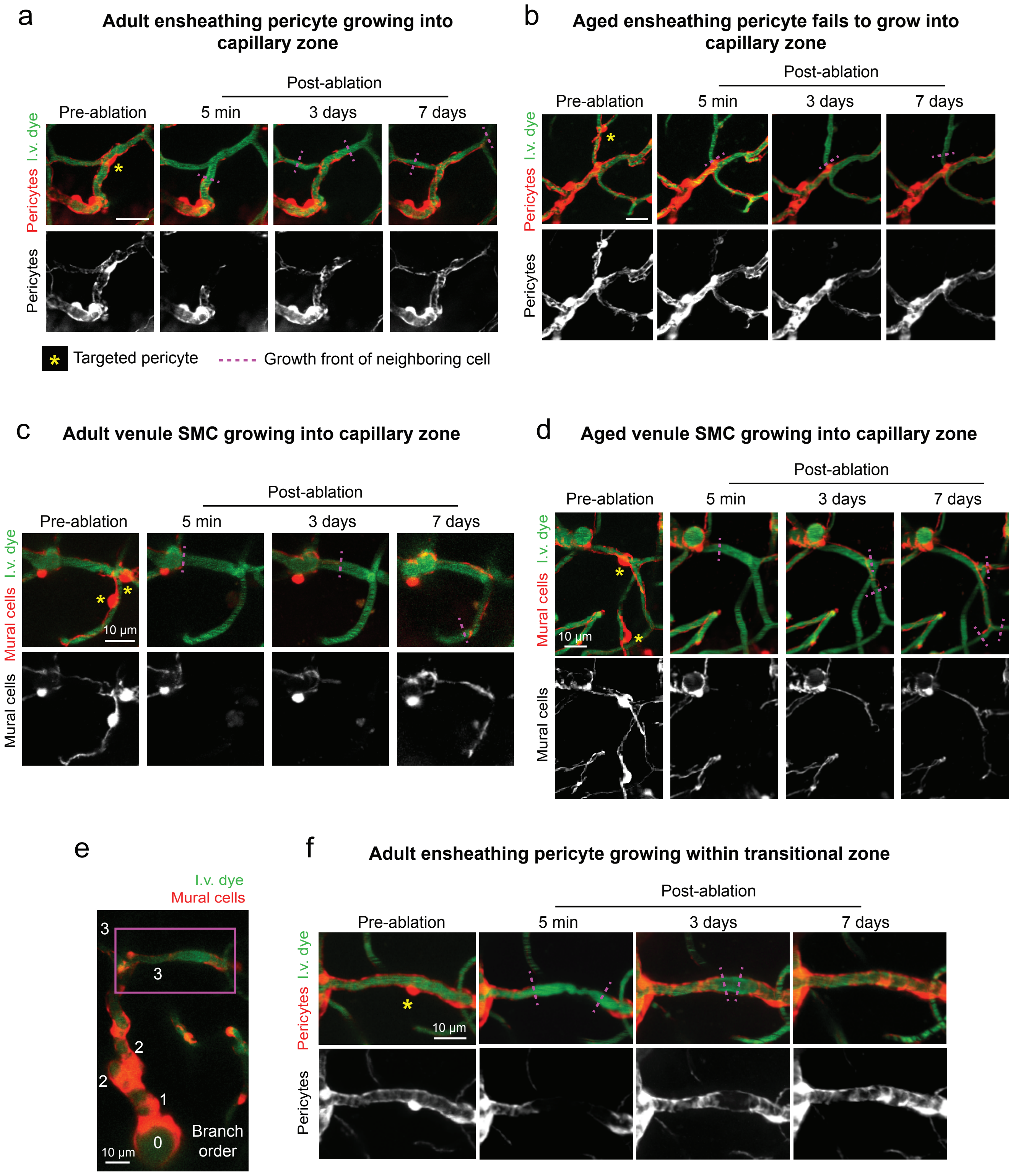

**Supplementary Fig. 5. Pericyte growth across microvascular zones in the adult and aged brain. (a)** An ensheathing pericyte from an adult mouse remodeling into the capillary zone. Ablated cells marked with a yellow asterisk, and growth front of processes marked with magenta dashed lines. Representative of 13 remodeling ensheathing pericytes in adult mice. I.v. dye = intravenous dye. **(b)** An ensheathing pericyte failing to remodel into the capillary zone over 7 days in an aged mouse. Representative of 12 ensheathing pericytes in aged mice. **(c)** Venule SMC from an adult brain growing into the capillary zone. Representative of 7 vSMCs from adult mice. **(d)** Venule SMC extending onto a capillary in an aged brain. Representative of 11 vSMCs from aged mice. **(e)** Image of an arteriole-capillary transition zone, with branch orders from the penetrating arteriole (0) marked. **(f)** Inset from (e), showing a 3rd order vessel with an ensheathing pericyte targeted for ablation. Over the next 7 days, neighboring ensheathing pericytes extend their processes into the area and their processes maintain an ensheathing morphology. Representative of 3 ensheathing pericyte ablations.

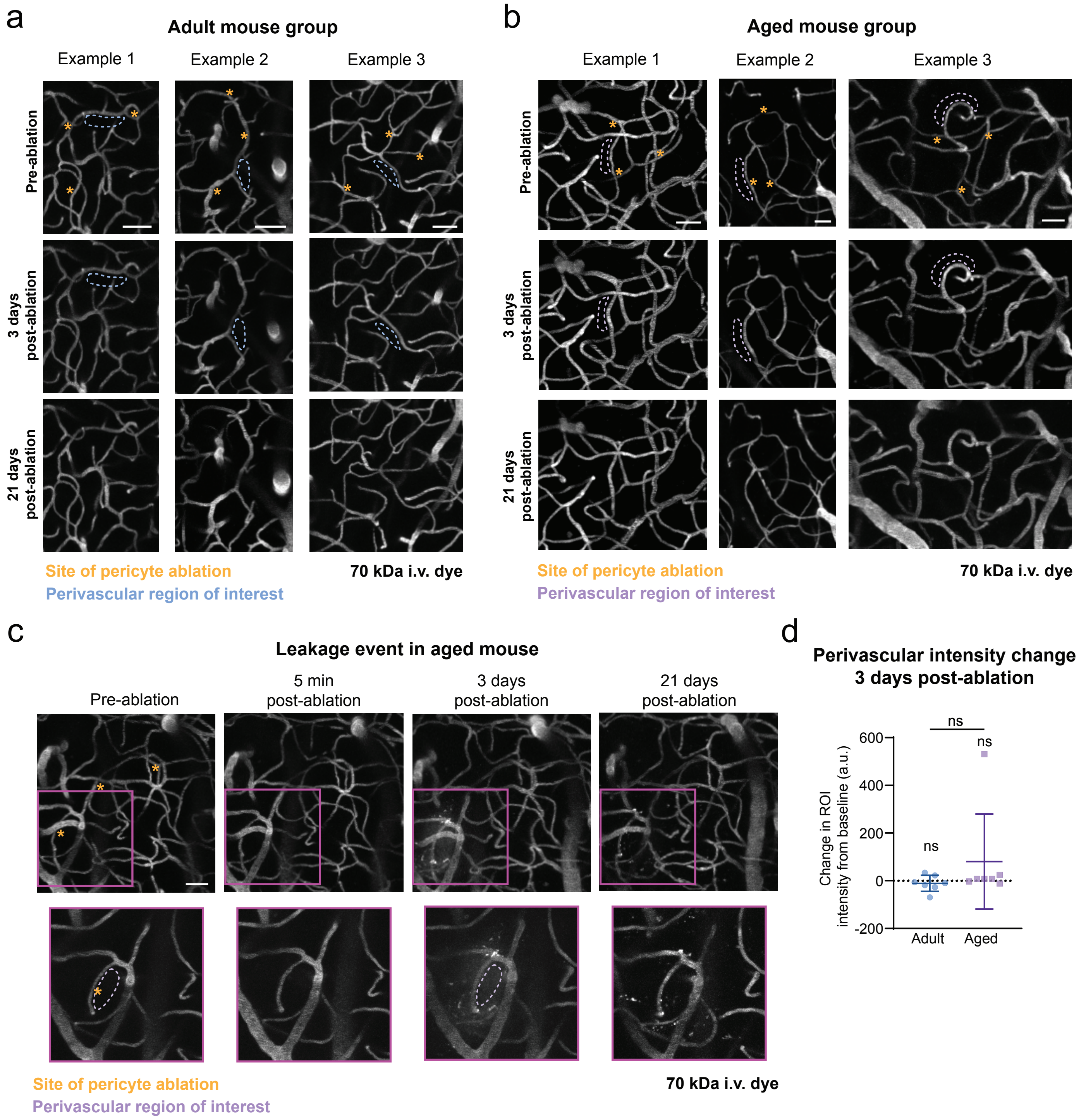

**Supplementary Fig. 6. Blood-brain barrier leakage is very rare with focal loss of pericyte coverage. (a)** Three examples of vasculature before and at two time points after triple pericyte ablation in adult mice. Sites of ablation are denoted with yellow asterisks, and perivascular regions of interest (ROI) for quantification of i.v. dye extravasation are in blue dashed lines. **(b)** Three examples of the vascular channel before after triple pericyte ablation in aged mice. Asterisks indicate location of pericyte ablation and ROIs for leakage assessment in dashed lines. Note the lack of 70 kDa intravenous (i.v.) dye permeating outside of the vessel walls across all examples. **(c)** (Top) A singular case of active leakage of 70 kDa i.v. dye noted at 3 days post-ablation in an aged mouse. No leakage occurred at 5 minutes post-ablation, suggesting that this was not caused by acute laser damage. (Bottom) Insets showing high resolution images of the capillary segment of interest. Sites of ablation are marked with asterisks, and dash ROI shows where parenchymal signal intensity measurements were taken. Scale bars = 50 µm. **(d)** Quantification of i.v. dye extravasation from perivascular locations lacking pericyte coverage at 3 days post ablation compared to pre-ablation. No parenchymal intensity change detected after pericyte loss in adult group, T(6)=0.8406, p=0.4328, by one-sample t test (two-sided). No perivascular leakage detected on average in aged group, T(6)=1.070, p=0.3257 by one-sample t test (two-sided). Singular example of leakage in this group can be seen as outlier driving large SD. No difference in intensity change between groups, T(6.345)=1.196, p=0.2746, by t test with Welch's correction for unequal variances (two-sided). N=7 ablation regions from 6 adult mice, and n=7 ablation regions from 6 aged mice. Data shown as mean ± SD.

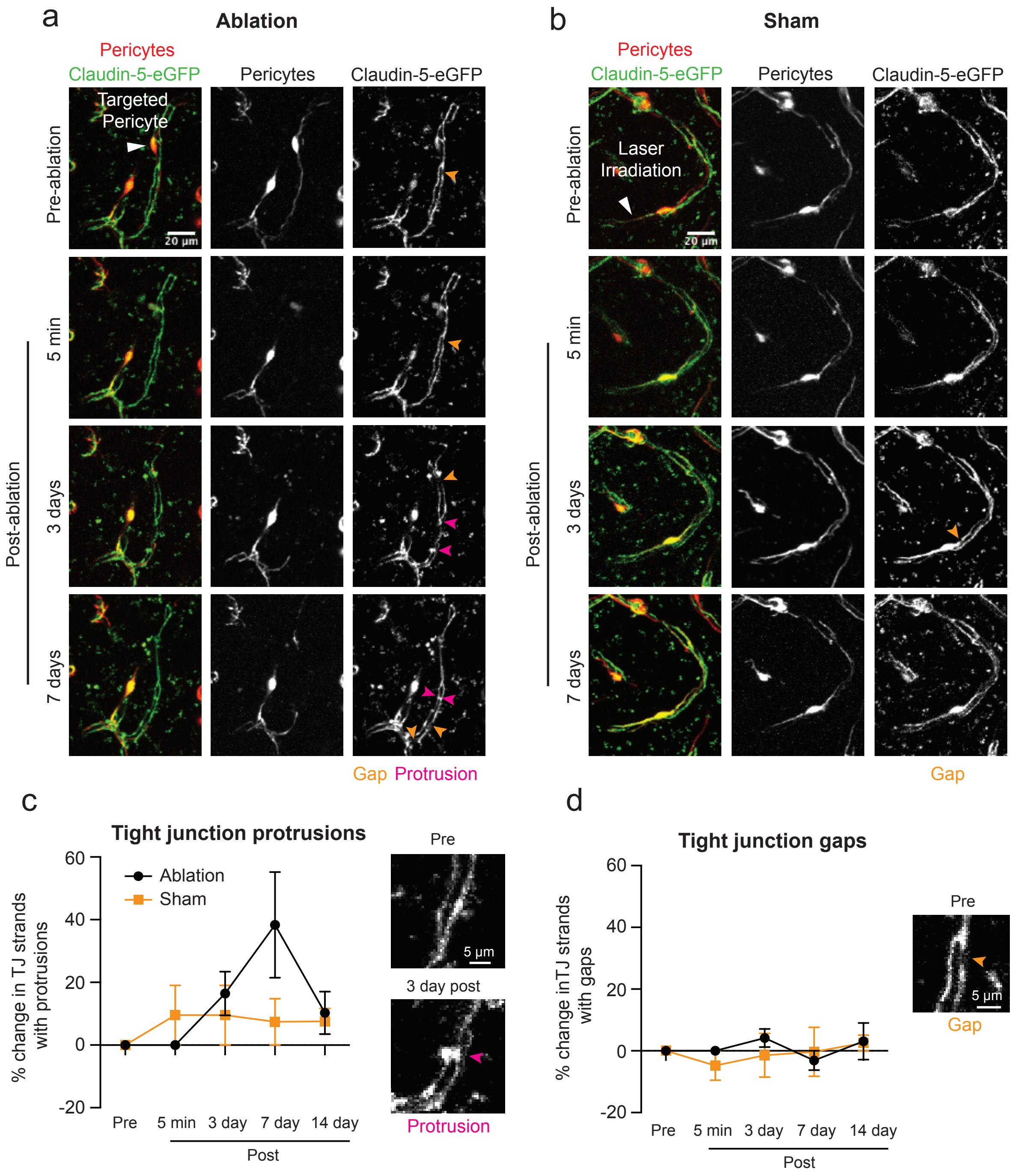

**Supplementary Fig. 7. No significant change in tight junction structure following the loss of individual pericytes. (a,b)** In vivo two-photon images from adult Pdgfrb-tdtomato;Claudin5-eGFP mice, showing pericytes and tight junction (TJ) structure before and 5 min, 3 days, and 7 days following (a) pericyte ablation or (b) off-target sham irradiation. Orange arrowheads denote gaps in the TJ strands, and pink arrowheads denote protrusions. **(c)** No significant difference in the percentage of TJ strands containing protrusions between the ablation and off-target sham irradiation groups at any post-irradiation timepoint compared to baseline. However, at day 7 there is a trend towards an increased number of protrusions in the ablation group compared to the control group. Two-way repeated measures ANOVA; no interaction detected between time and experiment type, $F(4,20)=2.317$, $p=0.0925$. **(d)** No significant difference in the percentage of TJ strands containing gaps between the ablation and control groups at any post-irradiation timepoint compared to baseline. Two-way repeated measures ANOVA; no interaction detected between time and experiment type, $F(4,20)=0.5185$, $p=0.7231$. For (c) and (d), n=4 pericyte ablations and 3 control irradiations (2 off-target irradiations and 1 failed pericyte ablation) from 3 mice aged 7-12 months. Data shown as mean ± SEM.

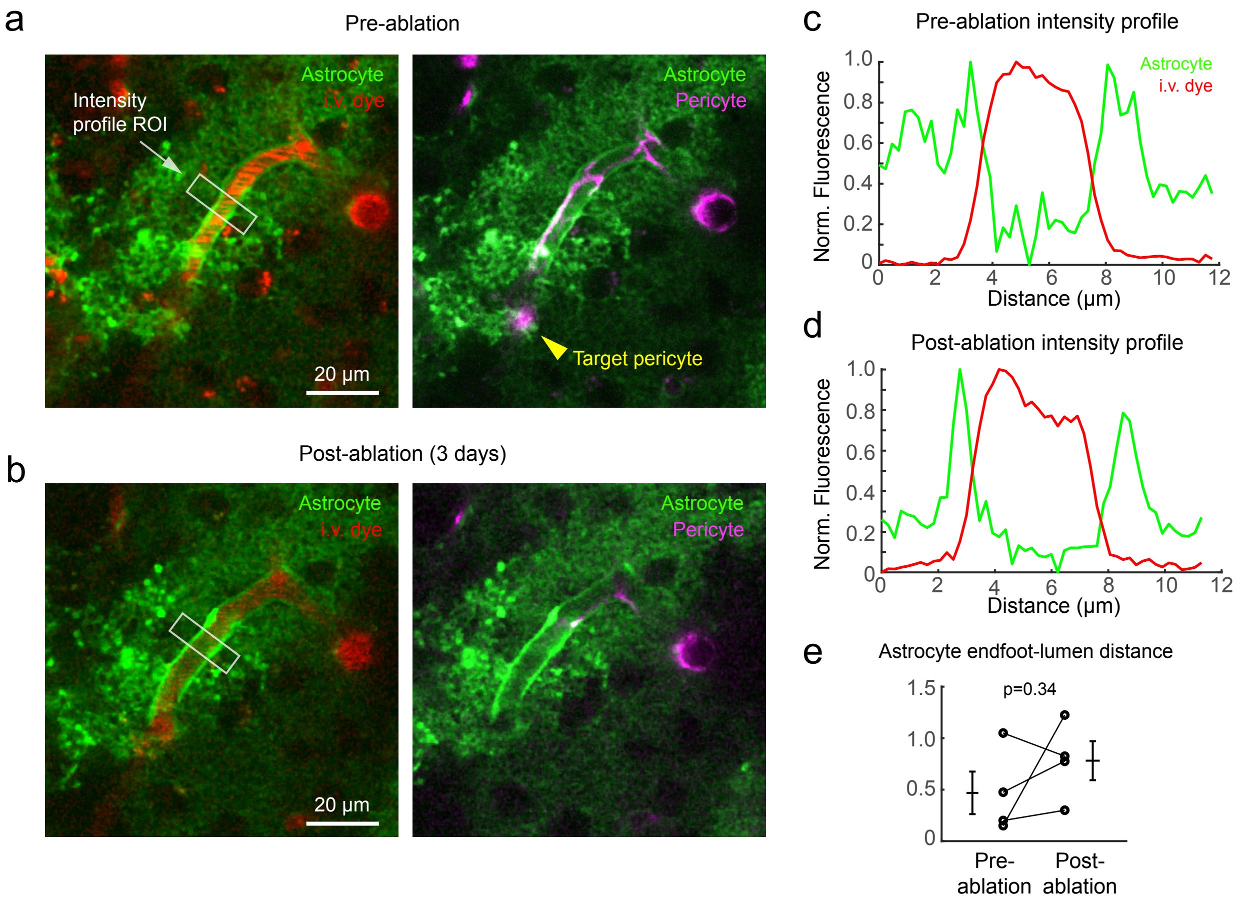

**Supplementary Fig. 8. Apposition of astrocyte endfeet to the capillary wall is not overtly disrupted by focal pericyte loss. (a,b)** In vivo two-photon images from aged Pdgfrb-tdtomato mice expressing Lck-GFP in astrocytes, before and 3 days after pericyte ablation. Left images show astrocytes (GFP) together with intravenous (i.v.) dye (Alexa 680-dextran) and right images show astrocyte together with pericytes. Here, a single pericyte has been ablated and the fluorescence intensity of astrocyte and i.v. dye signal is taken across a region affected by the pericyte ablation (white square). **c,d)** Normalized fluorescence intensity of astrocyte and i.v. dye signals plotted as a function of distance across the capillary cross-section at pre-ablation and post-ablation time-points. Note how sharp GFP peaks flanking the capillary lumen are still present after pericyte loss, suggesting intact endfoot coverage. **(e)** The distance between the intravascular wall (measured by FWHM) and the peak of the GFP signal does not change with pericyte loss. N=4 separate pericyte ablations were performed in 2 aged mice. Repeated-measures t-test (two-sided), t(3)=1.131, p=0.3402. Data shown as mean ± SEM.

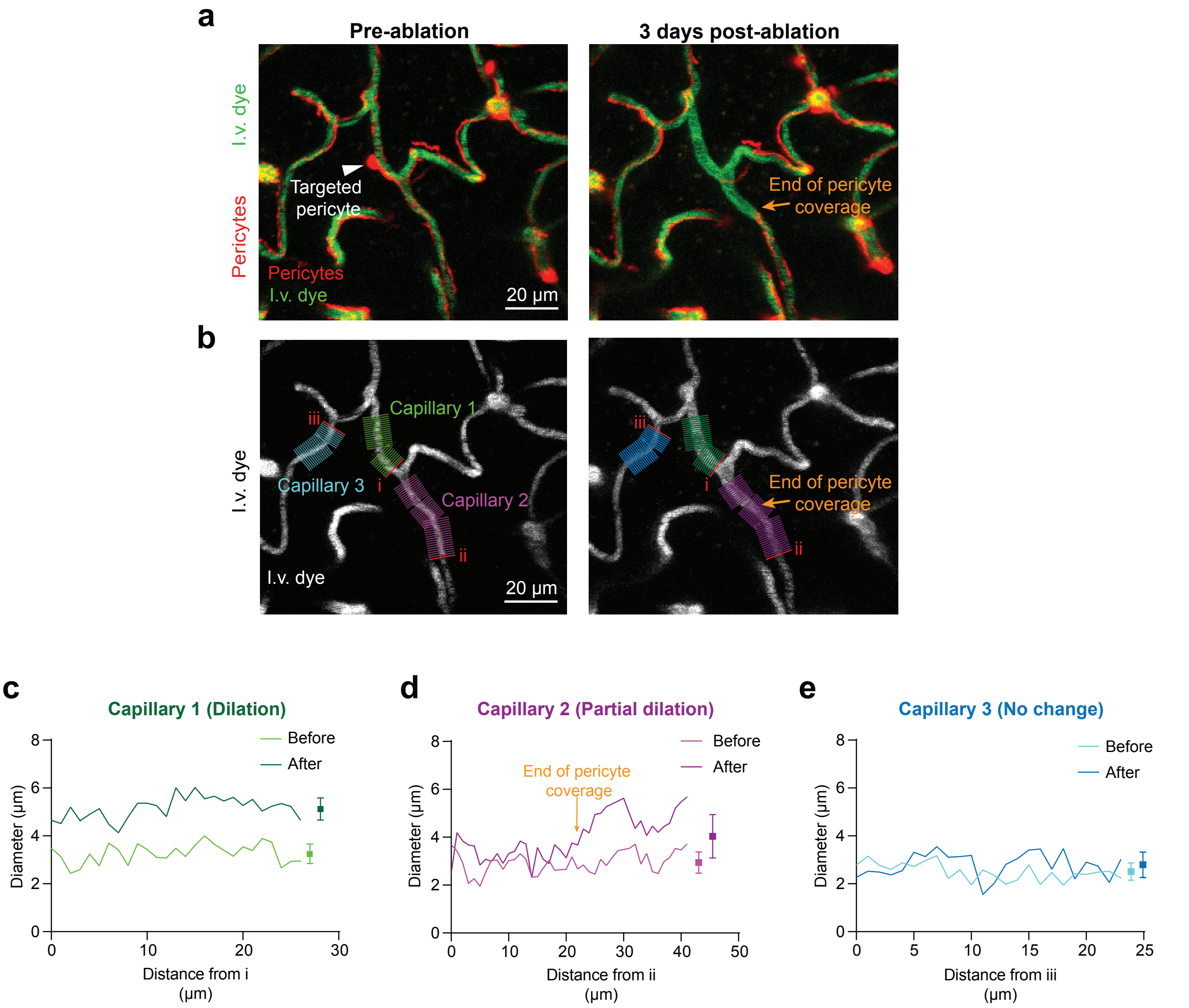

**Supplementary Fig. 9. Dilations are localized to portions of the capillary segment lacking pericyte coverage. (a)** Example of a capillary region before and 3 days after a pericyte ablation in an 18 month-old mouse. The edge of new process growth is marked with an orange arrow at 3 days post-ablation. I.v. dye = intravenous dye. **(b)** Corresponding i.v. dye channel, showing crosslines at which full-width at half-maximum intensity measurements were taken for diameter. Three individual capillary segments are examined. The edge of new pericyte process growth along Capillary 2 is marked at 3 days post-ablation. **(c)** Diameter of a fully uncovered capillary at Day 3 shows consistent dilations across its entire length. **(d)** A partially re-covered segment is dilated only beyond the end of a remodeling process. **(e)** A capillary segment not uncovered by ablation shows no change in diameter at 3 days compared to its baseline diameter. Data shown as mean ± SEM.

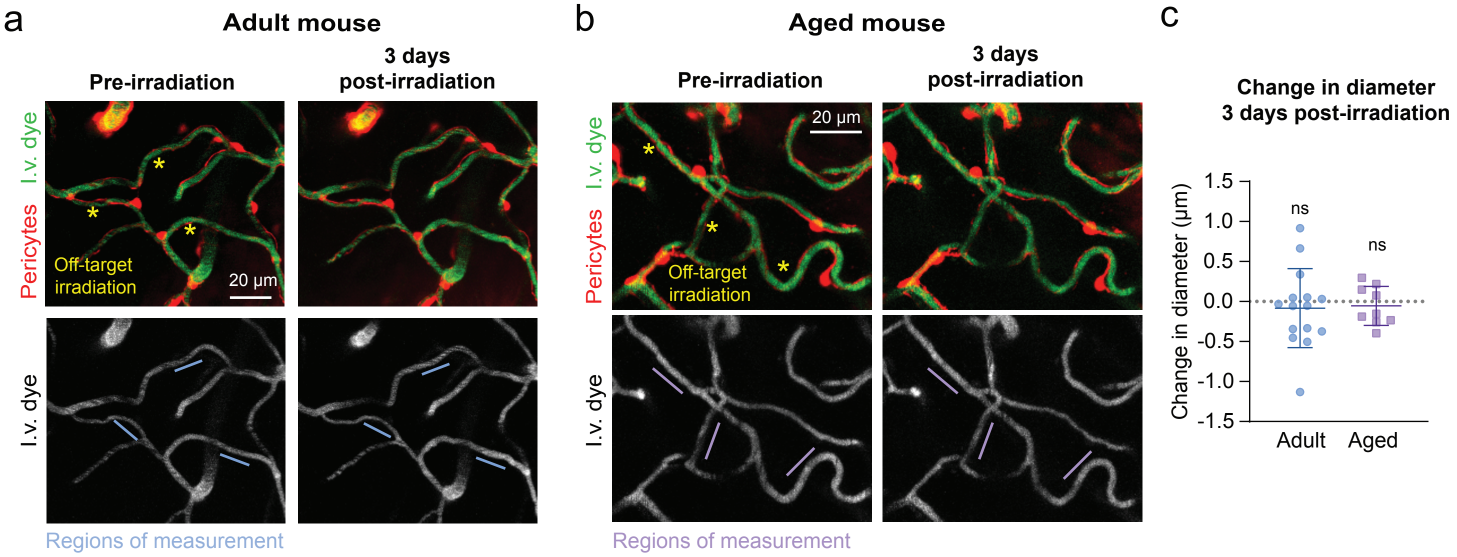

**Supplementary Fig. 10. No dilations detected 3 days following off-target sham irradiation. (a)** Example of a triple off-target irradiation in an adult mouse. Areas of irradiation are marked with yellow asterisks. Capillary diameters collected over time at segments marked with blue lines. I.v. dye = intravenous dye. **(b)** Example of a triple off-target irradiation in an aged mouse, as shown with yellow asterisks. Purple lines show segments measured for diameter. **(c)** No change in diameter in either age group at 3 days post-irradiation compared to baseline. For adults, t(14)=0.6475, p=0.5278 for 15 capillaries from 4 adult mice by one-sample t test (two-sided). For aged, t(8)=0.6732, p=0.5198 for 9 capillaries from 3 aged mice by one-sample t test (two-sided). Data shown as mean ± SD.

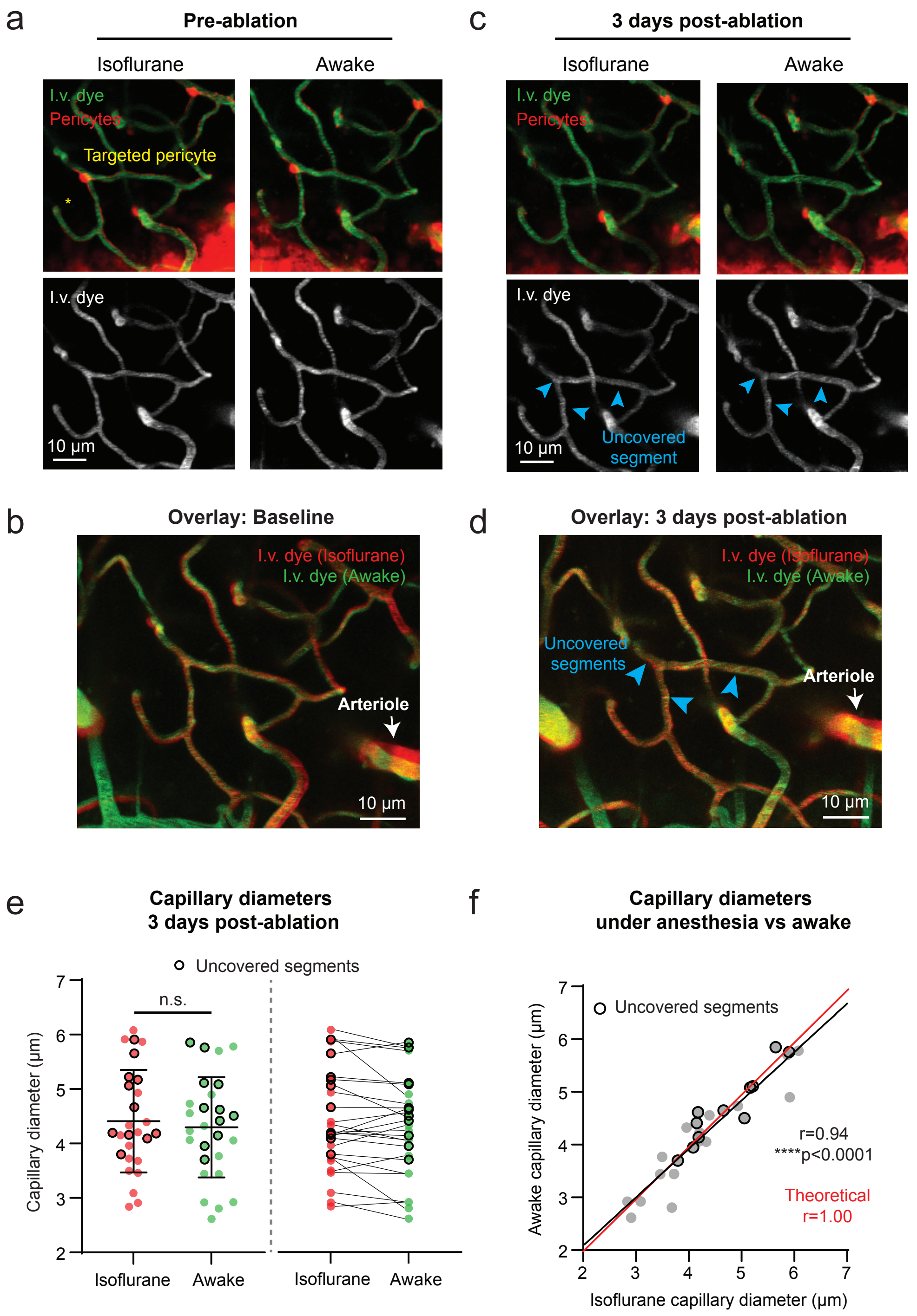

**Supplementary Fig. 11. Comparison of capillary diameters in awake versus isoflurane anesthesia conditions in aged mice. (a)** In vivo two-photon images of the same capillaries from an aged mouse, taken while under (left) isoflurane anesthesia (1.5% MAC) and (right) awake state. Pericyte targeted for ablation is marked with an asterisk. I.v. dye = intravenous dye. **(b)** Overlay of capillary area from (a), with image from anesthetized state in red and image from awake state in green. Note the presence of an artery within the field of view, which dilates under isoflurane and therefore serves as a positive control. **(c)** Capillary images 3 days post-ablation of the target pericyte, under both isoflurane and awake imaging conditions. Capillary segments lacking pericyte coverage are marked by arrowheads. **(d)** Overlay of capillary images obtained under isoflurane or awake 3 days post-ablation. **(e)** (Left) Quantification of capillary diameters in aged mice having undergone single pericyte ablation 3 days prior. Paired t-test (two-sided), $t(26) = 1.819$, $p=0.0804$. Diameters from uncovered segments are circled in black. N=27 capillaries (11 uncovered segments) in 3 separate ablation regions from 2 aged mice (1 female) . (Right) Scatter plot of same data, showing diameter change in individual capillaries going from anesthetized to awake state. Data shown as mean ± SD. **(f)** Correlation of diameters in isoflurane vs awake 3 days post-ablation, with segments uncovered and dilated circled in black. Pearson correlation (two-sided), $r=0.9382$, ****$p<0.0001$. Theoretical perfect correlation is shown as a red line for comparison.

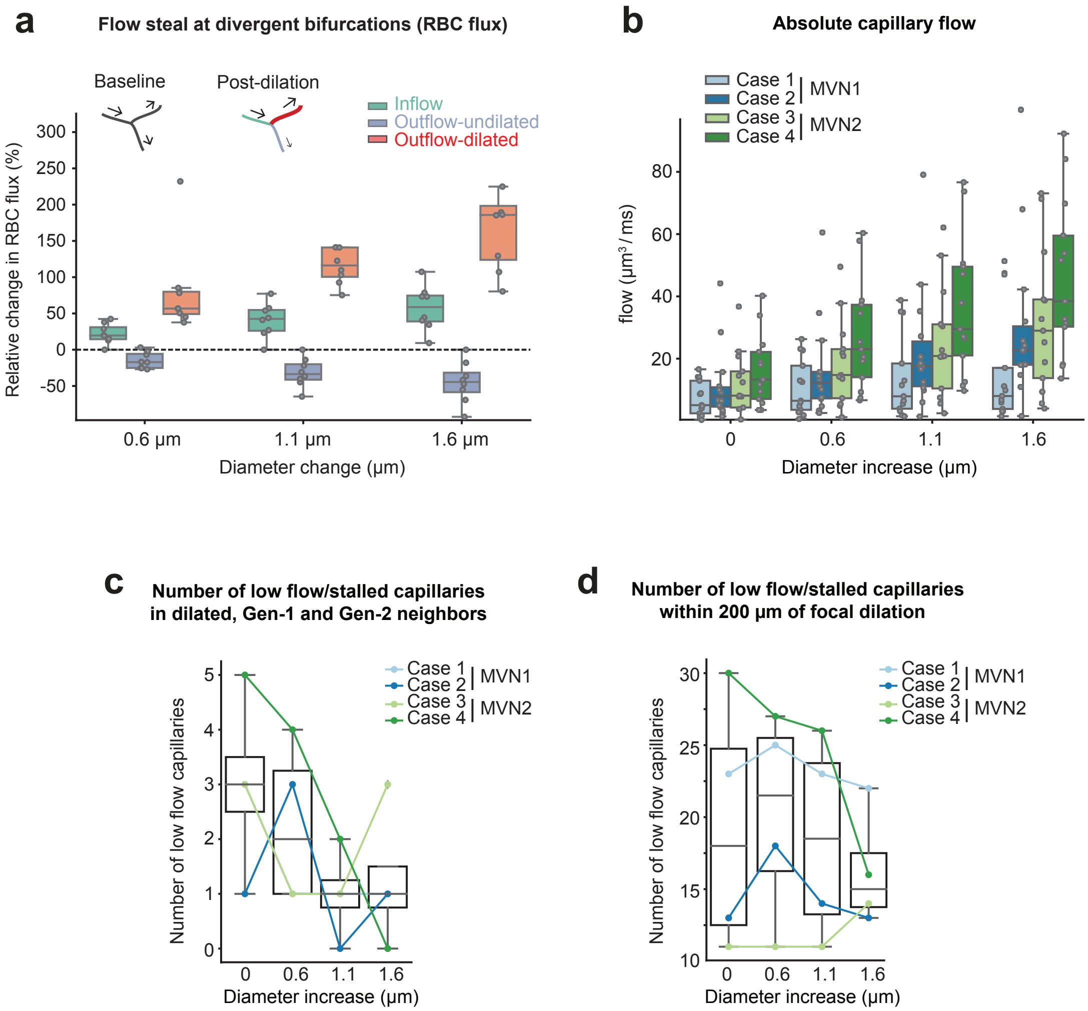

**Supplementary Fig. 12. Capillary flow changes after focal capillary dilation in silico. (a)** Flow steal at divergent bifurcations with one dilated outflow capillary shown with change in red blood cell (RBC) flux. N(0.6,1.1,1.6) = 24 vessels each. **(b)** Flow rate in dilated vessels for baseline (diameter increase = 0) and with increasing extent of dilation (diameter increase 0.6 μm, 1.1 μm and 1.6 μm). Number of dilated vessels: 13-14 (see Supplementary Table 1). Case 2 and Case 3 are in the center of the capillary bed, whereas Case 1 is closer to the venule side and Case 4 is closer to the arteriole side. N(0.0,0.6,1.1,1.6) = 53 vessels each, combined across cases. MVN = microvascular network. **(c,d)** Number of low flow or stalled capillaries in different vessel categories (c) or within a distance of 200 μm from the center of dilated vessels (c) does not increase with greater levels of focal capillary dilation. The threshold for low flow capillaries for the in silico study is defined as the lowest 5% of the capillary flow rates in the upper 200 μm of the microvascular networks. N(0.0,0.6,1.1,1.6) = 4 MVN cases. For all box and whisker plots, center = median, box bounds = upper (Q3) and lower (Q1) quartiles, whiskers = last data point within Q1 - 1.5*(Q3-Q1) and Q3 + 1.5*(Q3-Q1).

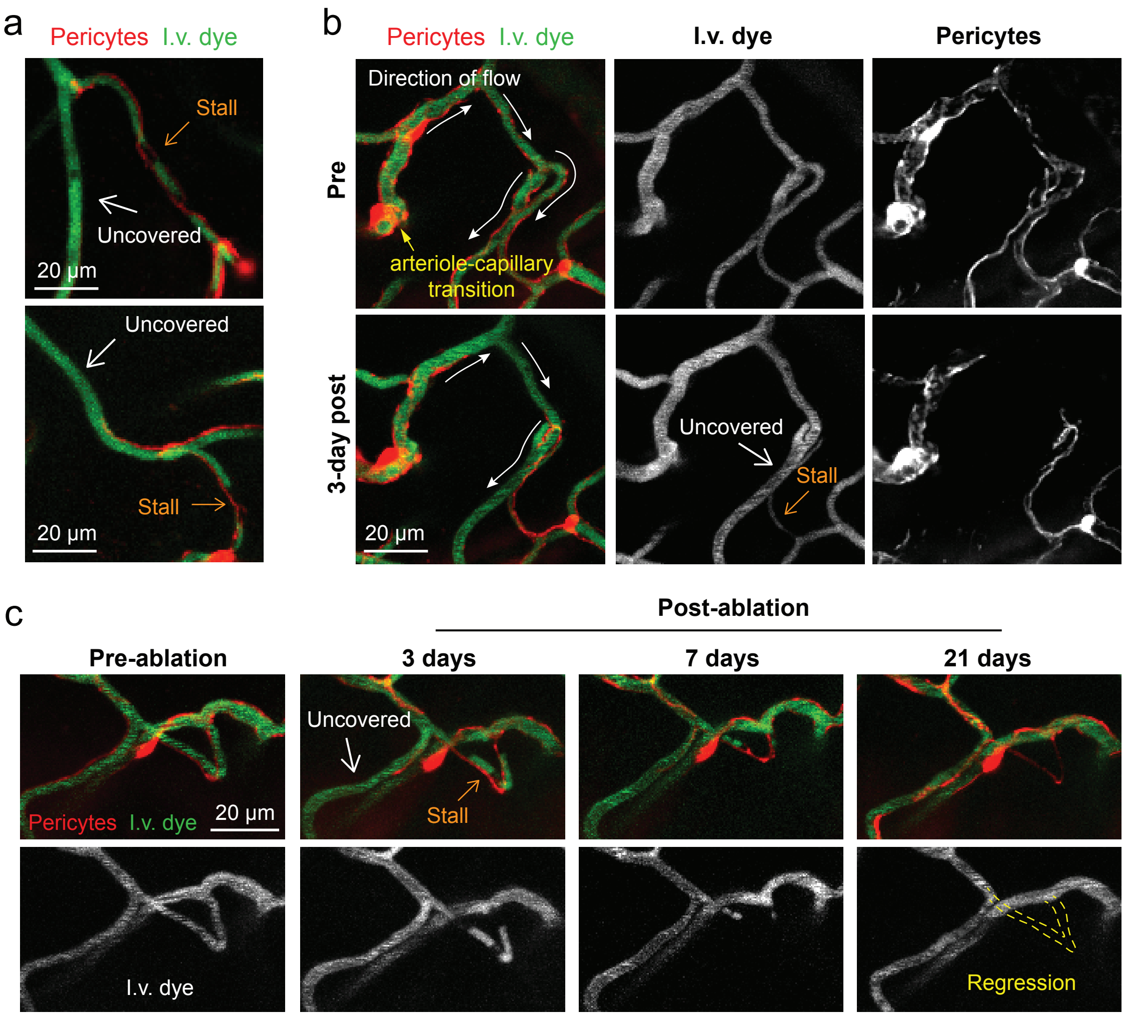

**Supplementary Fig. 13. Additional examples of capillary flow stalls and regressions. (a)** Two additional images from aged mice of blood flow stalls in capillary segments immediately adjacent to capillaries uncovered by pericyte ablation. I.v. dye = intravenous dye. **(b)** Detailed view of blood flow redirection (arrows) in capillaries downstream from the arteriole-capillary transition zone. The pericyte loss-induced dilation in one segment 3 days post-ablation results in the complete steal of flow away from the alternate, pericyte-covered segment. Example from an adult mouse. **(c)** The development of a capillary regression in an adult mouse over time following prolonged capillary stalling. A blood flow stall is observed 3 days post-ablation, in a capillary adjacent to a pericyte-uncovered, dilated segment. This capillary is undergoing regression by 7 days post-ablation, a process which is fully complete by 21 days post-ablation. All images representative of stalls/regression observed across 14 pericyte ablation experiments.

**Supplementary Tables. Berthiaume *et al.***

**Supplementary Table 1. Overview of selection criteria to choose potential *base capillaries* around which pericyte ablation will be mimicked.**

| | Cortical Depth [μm] | Distance to main branch [branches] | | Diameter [μm] | RBC flux [RBCs/s] | RBC velocity [mm/s] | Distance to center MVN1/ MVN2 [μm] | Distance to boundary [branches] |
|---|---|---|---|---|---|---|---|---|
| | | DA | AV | | | | | |
| Min | 20 | 4 | 1 | 2.17 | 17 | 0.19 | - | 2 |
| Max | 120 | - | - | 4.81 | 381 | 2.70 | 319 / 350 | - |

DA: descending arteriole. AV: ascending venule.

**Supplementary Table 2. Baseline characteristics for the set of affected capillaries around different base capillaries.**

| MVN | No. affected caps. [-] | Cortical depth [μm] | Diameter [μm] | RBC flux [cells/s] | RBC velocity [mm/s] | Length [μm] | Total length [μm] | Bounding box ΔX /ΔY/ΔZ [μm] |
|---|---|---|---|---|---|---|---|---|
| 1 | 13 | 47 ± 36 | 3.4 ± 0.4 | 44 ± 32 | 1.0 ± 0.5 | 67 ± 44 | 870 | 93/119/150 |
| 1 | 14 | 98 ± 64 | 3.5 ± 0.5 | 26 ± 21 | 0.6 ± 0.3 | 66 ± 50 | 926 | 160/158/301 |
| 1 | 13 | 70 ± 49 | 3.7 ± 0.5 | 39 ± 34 | 0.7 ± 0.6 | 59 ± 41 | 768 | 90/150/148 |
| 1 | 13 | 42 ± 22 | 3.6 ± 0.7 | 29 ± 21 | 0.4 ± 0.3 | 48 ± 33 | 618 | 62/260/68 |
| 1 | 13 | 115 ± 54 | 4.3 ± 0.7 | 14 ± 12 | 0.4 ± 0.3 | 105 ± 91 | 1365 | 193/238/232 |
| 1 | 14 | 89 ± 35 | 3.6 ± 0.5 | 59 ± 56 | 1.2 ± 0.9 | 60 ± 24 | 836 | 150/145/140 |
| 1 | 13 | 146 ± 67 | 3.7 ± 0.4 | 36 ± 35 | 0.9 ± 0.7 | 83 ± 59 | 1081 | 134/179/212 |
| 2 | 15 | 152 ± 59 | 3.8 ± 0.7 | 73 ± 54 | 0.9 ± 0.5 | 68 ± 38 | 1018 | 148/85/229 |
| 2 | 17 | 140 ± 53 | 3.7 ± 0.5 | 93 ± 104 | 1.4 ± 1.0 | 60 ± 33 | 1021 | 90/123/209 |
| 2 | 15 | 41 ± 47 | 4.2 ± 0.7 | 230 ± 171 | 1.8 ± 0.9 | 55 ± 53 | 823 | 102/67/265 |
| 2 | 13 | 93 ± 33 | 3.7 + 0.6 | 115 ± 104 | 1.6 ± 0.8 | 64 ± 32 | 829 | 154/184/142 |
| 2 | 13 | 73 ± 20 | 3.8 ± 0.7 | 187 ± 141 | 2.0 ± 1.2 | 47 ± 24 | 616 | 106/200/52 |
| 2 | 13 | 124 ± 30 | 3.5 ± 0.6 | 121 ± 101 | 1.2 ± 0.7 | 58. ± 41 | 755 | 155/146/142 |
| 2 | 13 | 76 ± 37 | 3.9 ± 1.1 | 62 ± 34 | 0.9 ± 0.5 | 52 ± 30 | 673 | 102/106/158 |

For cortical depth, diameter, RBC flux and RBC velocity the mean ± SD across all affected capillaries is provided. MVN: Microvascular network. No. affected caps.: Number of affected capillaries. Bounding box: dimensions of a bounding box around all affected capillaries. Cases chosen for further analyses are highlighted in gray.
